# Reciprocal Regulation Between the SCF^FBXO24^ Ubiquitin E3 Ligase and FoxP1 Protein

**DOI:** 10.1101/2025.05.23.653845

**Authors:** Abigail Maloy, Sydney Walter, Arthur Mascilli, David C. Klein, Santana M. Lardo, James Londino, Toru Nyunoya, John McDyer, Mia Sun, Xuemei Zeng, Nathan Yates, Pamela Cantrell, Sarah J. Hainer, Rama K. Mallampalli, Divay Chandra

## Abstract

Forkhead Box Protein P1 (FoxP1) is a crucial transcriptional repressor essential for the development of the brain and heart. In adults, FoxP1 protein levels are dysregulated in a variety of disorders, including chronic obstructive pulmonary disease (COPD), atherosclerosis, and heart failure, where they causally contribute to disease pathogenesis. Although independent investigators have reported that FoxP1 protein is ubiquitinated, and E3 ligases have been identified for other FoxP family proteins, the identity of the E3 ligase that controls FoxP1 protein stability has remained unknown. Here, we identify FBXO24, a subunit of the Skp-Cullin-F-box (SCF) ubiquitin E3 ligase complex, as the regulator of FoxP1 ubiquitination and stability. Specifically, FBXO24 regulates K48 and K63 ubiquitination, complexes with, and co-localizes to the nucleus with FoxP1 protein in lung epithelial cells. Depleting FBXO24 reverses the unfolded protein response and cell death triggered by loss of FoxP1 protein in lung epithelium, suggesting a protective role. Additionally, FBXO24 knockout mice exhibit elevated FoxP1 levels in the lung and heart and reduced unfolded protein response activity after short-term cigarette smoke exposure. Intriguingly, we also uncovered bidirectional regulation, whereby FoxP1 protein binds to the FBXO24 promoter to suppress FBXO24 transcription. To our knowledge, this is the first evidence that a substrate for an E3 ligase can also regulate the E3 ligase and, therefore, control levels of other substrates, revealing new regulatory networks. Targeting FBXO24 may offer a therapeutic strategy for COPD, atherosclerosis, and heart failure by stabilizing FoxP1 levels in the heart and lungs and mitigating harmful downstream effects.

## INTRODUCTION

The Forkhead box (Fox) family includes transcription factors in subclasses A-S characterized by a forkhead box DNA binding domain (1). These proteins are conserved from yeast to mammals and regulate diverse biological processes during development and adult life (1, 2). Four subclass P proteins exist in mammals (FoxP1-4) (3). FoxP1 protein contains 677 amino acids and is 93% conserved between humans and mice.

FoxP1 is essential for heart and brain development (1). Specifically, FoxP1^-/-^ mice die by embryonic day 14 due to cardiac malformations (4). Also, mutations or deletions in the FoxP1 gene cause a neurodevelopmental disorder in humans termed *FoxP1 Syndrome*, characterized by intellectual disability, language deficits, autism spectrum disorder, hypotonia, and congenital abnormalities (5).

Recent studies have found that FoxP1 is critical not only for organ development but also to counteract prevalent cardiovascular and pulmonary diseases in adult life, such as atherosclerosis, heart failure, and COPD. Specifically, FoxP1 is a critical anti-atherosclerotic mediator and gatekeeper of vessel inflammation. It counteracts vessel inflammation by suppressing endothelial inflammasome components Nlrp3, caspase-1, and IL-1β (6). In human coronary atherosclerotic plaques and atherosusciptable endothelium, FoxP1 protein levels are reduced, leading to increased vessel inflammation (6). In agreement, increasing FoxP1 protein levels in murine coronary endothelium suppresses atherosclerosis (6).

In terms of heart failure, FoxP1 protein counteracts cardiac fibrosis and hypertrophy, i.e., the precursors of cardiac failure (7–9). Specifically, FoxP1 directly suppresses TGF-β1 in the cardiac endothelium, thereby inhibiting myofibroblast formation and extracellular matrix protein production (9). FoxP1 protein levels are significantly downregulated in the cardiac endothelium during angiotensin II-induced cardiac remodeling, and restoring FoxP1 reverses cardiac remodeling and dysfunction.

In lung disease, the FoxP1 gene has emerged as a novel COPD susceptibility locus in genome-wide association studies (10). Our previous work demonstrated that FoxP1 protein is depleted in the lungs of individuals with COPD (11). Functionally, FoxP1 binds to and represses the promoters of key genes involved in the unfolded protein response (UPR) within the lung epithelium (11). Knockdown of FoxP1 in lung epithelial cells led to dysregulated activation of the UPR, increased production of matrix metalloproteinases and inflammatory mediators, and enhanced cigarette smoke-induced emphysema *in vivo* (10). These findings suggest a critical role for FoxP1 in maintaining lung epithelial homeostasis and preventing pathological lung remodeling in response to environmental insults.

Therefore, loss of FoxP1 protein is a key pathogenic mechanism in prevalent diseases such as atherosclerosis, heart failure, and COPD and restoring FoxP1 protein levels may be a viable therapeutic strategy for these disorders. FoxP1 protein levels can be augmented by inhibiting its ubiquitination.

Ubiquitination controls the lifespan of most cellular proteins and involves a hierarchical, exquisite system. Ubiquitination involves the stepwise transfer of ubiquitin from an E1 ubiquitin-activating enzyme to an E2 ubiquitin-conjugating enzyme and finally to an E3 ubiquitin ligase complex (12, 13). In the last step of the reaction, the E3 ubiquitin ligase transfers ubiquitin chains to the substrate to facilitate degradation by the 26S proteasome or sorting to the endosome-lysosome pathway. Two E1 enzymes, almost 40 E2 enzymes, and more than 1000 E3 enzymes have been identified in mammalian cells (12). The Skp-Cullin-F-box (SCF) superfamily represents the largest group of E3 ligases that mediate critical roles in cell biology, including tumorigenesis and inflammation. Within the SCF apparatus is a receptor module, termed F-box protein, that binds substrates.

Although independent investigators have reported that FoxP1 protein is ubiquitinated (14), and E3 ligases have been identified for other FoxP proteins (14), the identity of an E3 ligase that controls FoxP1 protein ubiquitination and stability has remained unknown.

Here, we identified that a subunit of the Skp-Cullin-F-box ubiquitin E3 ligase apparatus, i.e., FBXO24, controls ubiquitination and the stability of FoxP1. Specifically, FBXO24 regulates K48 and K63 ubiquitination, complexes with and co-localizes to the nucleus with FoxP1 in lung epithelial cells. Further, depleting FBXO24 reverses the increase in UPR activity and cell death induced by the knockdown of FoxP1 protein in the lung epithelium. Also, FBXO24^-/-^ mice demonstrate increased FoxP1 protein levels in the heart and the lung and less UPR activity in the lung after exposure to cigarette smoke for two weeks. Therefore, inhibiting FBXO24 may be a therapeutic strategy for atherosclerosis, heart failure, and COPD as it leads to higher FoxP1 protein levels and counteract deleterious downstream pathways. Finally, we identified bidirectional regulation whereby FoxP1 binds to the promoter of FBXO24 and regulates FBXO24 mRNA levels. To our knowledge, this is the first evidence that a substrate for an E3 ligase can also regulate the E3 ligase and, therefore, control levels of other substrates, revealing new regulatory networks.

## MATERIALS AND METHODS

### Cell culture and transfection

Murine lung epithelial (MLE12) cells (ATCC) and human bronchial epithelial (BEAS-2B) cells (ATCC) were cultured with HITES medium (DMEM/F12 supplemented with insulin, transferrin, hydrocortisone, β-estradiol, and glutamine) containing 10% FBS and antibiotics as described previously (15).

For knockdown, small interfering RNAs (siRNA) were purchased from Integrated DNA Technologies (Coralville, Iowa), along with control/nonsense dsiRNA. The cell culture medium was changed when cells were 70-80% confluent. Scramble or targeted dsiRNA (final concentration 50 nM) was mixed with transfection buffer and GenMute transfection reagent (SignaGen Labs, Rockville, MD) according to manufacturer instructions and added to respective wells. The medium was changed 6 h later, and lysate was collected 48 h later.

For overexpression, plasmids were transfected at the stated concentration with X-tremeGENE HP transfection reagent (Millipore, St. Louis, MO). FBXO24 cDNA sequence cloned into a pLX304 vector with a V5 tag (HsCD00446263), FBXO23 cDNA sequence cloned into a pLX304 vector with a V5 tag (HsCD00436300), and FoxP1 cDNA sequence cloned into a pLenti6.3 lenti vector with a V5 tag (HsCD00852556) were obtained from DNASU and were confirmed to contain the correct open reading frames by sequencing. Ub-K63-HA and Ub-K48-HA were a gift from Ted Dawson (Addgene plasmid # 17606 and #17605) (16).

### Immunoblotting

Supernatants were discarded, and the cells were washed twice with ice-cold PBS. Lysis buffer was added (0.5% Triton in PBS with 1x Halt protease inhibitor cocktail), and the lysate was mechanically scraped off and removed from the bottom of each well. Lysates were sonicated for 15 s on ice twice, protein concentration was measured (DC protein assay, Bio-Rad), and 15 μg/lane of lysate were loaded onto 10% stain-free SDS-PAGE gels (Bio-Rad).

After electrophoresis, the gels were photographed under UV light to image the protein in the gel according to the manufacturer’s instructions. After transfer to low fluorescence PVDF membranes (Millipore) using a Trans-Blot Turbo transfer system (Bio-Rad), the membranes were re-imaged under UV light to confirm the complete and even transfer of the protein from the gel. The membranes were then blocked with 3% (w/v) BSA in TBST (25 mM Tris-HCl, pH 7.4, 137 mM NaCl, and 0.1% Tween 20) for 1 h, and incubated with primary antibodies in 3% (w/v) BSA in TBST at 4 °C overnight. The membranes were then washed thrice with TBST at 5 min intervals followed by a 1 h incubation with horseradish peroxidase-conjugated secondary antibody (1:2000).

The membranes were developed with an enhanced chemiluminescence detection system according to the manufacturer’s instructions (Bio-Rad). Densitometry was performed and adjusted to total protein per lane using automated measurement and rolling disk normalization using Image Lab v5.2.1 according to manufacturer instructions (Bio-Rad). Total protein per lane was used to correct for protein loading rather than housekeeping genes (such as β-actin) because independent studies indicate that the total protein method is more accurate and reproducible (17–19).

### Immunoprecipitation

Once BEAS-2B cells were close to confluence in 10 cm dishes, they were treated with 25 μM MG132 for 1 h to inhibit the proteasome. Next, the supernatant was removed, and cells were washed with ice-cold PBS twice. RIPA buffer supplemented with 1x ubiquitin aldehyde (to inhibit deubiquitination) and 1x Halt protease inhibitor cocktail was added, and the lysate was mechanically scraped from the dishes.

Lysates were sonicated for 7 s on ice twice, protein concentration measured, and 500 μg protein was incubated with isotype control IgG antibody and protein A/G agarose to pre-clear. Samples were placed on a tube rotator at 4°C for 30 min and then centrifuged. The pellet was discarded, and the supernatant was incubated with 5 μg primary antibody or 5 μg isotype control IgG on a tube rotator at 4°C for 1 h. After 1 h, protein A/G agarose was added, and samples were incubated on a tube rotator at 4°C overnight. Samples were centrifuged, pellets were washed in PBS, and eluted in 1X Laemmli Buffer. The eluate was then subject to immunoblotting.

### RT-PCR

RNA was isolated from cells using RNeasy Mini Kits (Qiagen) per the manufacturer’s instructions. After their concentrations were measured, isolated RNAs were converted to cDNA using High-Capacity RNA-to-cDNA Kits (Life Technologies, Grand Island, NY). Real-time PCR assays were performed using SYBR® Select Master Mix for CFX (2X) (Life Technologies, Grand Island, NY) with the C1000 Thermal Cycler (BioRad, Hercules, CA) per manufacturer’s instructions. Target gene mRNA expression was normalized to that of a housekeeping gene (GAPDH).

### Cellular immunofluorescence

BEAS-2B cells were plated in 12-well plates with sterile coverslips placed at the bottom of each well. After the cells were transfected, coverslips were removed, the cells were washed thrice with PBS, fixed with 2% formaldehyde for 15 min, permeabilized with 0.1% Triton X 100 for 15 min, blocked with 2% BSA for 45 min, and incubated with 1:500 anti-V5 or anti-MYC antibody. After 1 h, the cells were washed, and 1:1000 goat anti-rabbit Cy5 secondary antibody was added. Finally, the cells were washed with PBS, mounted with DAPI containing mounting medium (Fluoroshield, Abcam), and imaged using a Nikon ECLIPSE A1 confocal microscope.

### Flow Cytometry

BEAS-2B cells were detached from culture dishes using dissociation buffer (0.5% BSA, 5 μM EDTA in PBS) for 20 mins at 4°C. The cell suspension was centrifuged, washed with 0.5% BSA once, and stained with LIVE/DEAD^™^ Fixable Aqua Dead Cell Stain Kit per manufacturer instructions (Invitrogen). After staining, cells were washed with 0.5% BSA and assessed using a BD LSRFortessa flow cytometer to capture at least 10,000 events per sample. Data was analyzed using FlowJo Software v10 (Ashland, OR).

### ChIP-Polymerase Chain Reaction (ChIP-PCR)

ChIP was performed using the Pierce Magnetic ChIP Kit (ThermoFisher Scientific, cat# 26157) per manufacturer’s instructions. In brief, BEAS-2B cells in culture dishes were incubated with formaldehyde (final concentration of 1%) for 10 mins to crosslink proteins to DNA. Next, the reaction was quenched by incubating with 10x Glycine solution (final concentration of 1x) for 5 min. The cells were then washed twice with ice-cold PBS and lysed by scraping in PBS supplemented with 1:100 protease inhibitor cocktail. Samples were centrifuged, and the pellet was resuspended in Membrane Extraction Buffer, vortexed, and incubated on ice for 10 min. Samples were then centrifuged, resuspended in MNase Digestion Buffer, and incubated with 1 μL of MNase at 37°C for 15 min to digest DNA. Next, MNase Stop Solution was added, and samples were sonicated for 20 sec 3 times at 20% power, with 20 sec breaks between sonications.

Input DNA was set aside, and the remaining DNA was incubated with anti-FoxP1 antibody (Abcam #16645) or isotype control antibody (normal rabbit IgG) overnight at 4°C on a rocker. The following day, samples were incubated with protein A/G Magnetic Beads for 2 h at 4°C on a rocker. The beads were then washed with IP Wash Buffer 1 & 2, three times each. Next, beads were eluted by incubating with IP Elution Buffer at 65°C for 40 min. IP eluate and input were then treated with NaCl and Proteinase K (final concentration of 0.2 M and 0.67 mg/mL, respectively) at 65°C for 90 min.

DNA was recovered by adding DNA Binding Buffer and incubating with a DNA Clean-Up Column that was washed with DNA Column Wash Buffer. DNA was eluted using DNA Column Elution Solution and used for PCR with DreamTaq polymerase with the primers that bind to the FBXO24 promoter region (see supplement for primers sequences). PCR product was run on a 1% agarose gel and imaged.

### Analysis of publicly available FoxP1 ChIP-seq data

We analyzed publicly available FoxP1 ChIP-seq data from human embryonic stem cells (GEO: GSE30992) and HepG2 cells (GEO: GSE105268), as data from pulmonary cells was not available (20). Sequencing reads (fastq files) were downloaded and mapped to the hg38 genome using Bowtie. HOMER was used to generate normalized UCSC browser tracks, which were visualized over the FBXO24 promoter (21).

### Prediction of FoxP1 binding sites

We matched the known DNA motif recognized by FoxP1(20) with the DNA sequence of the FBXO24 promoter using the “toPWM” command in the “TFBSTools” package in R (22).

### CUT&RUN

CUT&RUN was performed using the low salt elution method for fragment release, as described previously (23–25). Specifically, A549 cells were transfected with either FoxP1-V5 or vector backbone. After 48 h, nuclei were extracted using Nuclear Extraction (NE) buffer (20 mM HEPES-KOH, pH 7.9, 10 mM KCl, 0.5mM spermidine, 0.1% Triton X-100, 20% glycerol, & protease inhibitors) and flash frozen at -80°C for storage.

Nuclei were thawed on ice for 5 min, and 100,000 nuclei/condition were resuspended in 600 μL NE buffer. Next, nuclei were bound to pre-washed Concanavalin beads (Polysciences; 25 μL per 100,000 cells) in binding buffer (20 mM HEPES, pH 7.5, 150 mM NaCl, 0.5 mM spermidine, 0.1% BSA, 2 mM EDTA, & protease inhibitors) and washed with wash buffer (20 mM HEPES, pH 7.5, 150 mM NaCl, 0.5 mM spermidine, 0.1% BSA, & protease inhibitors). V5 antibody (Invitrogen cat# R960-25, 1:250) or control IgG (Sigma-Aldrich, cat# 06-371,1:250) were then added in 250 μL wash buffer for 1 h at RT. After two washes, pA-MNase was added to each sample in 250 μL wash buffer for 30 min at RT. After two more washes, samples were resuspended in 150 μL wash buffer, and 3 μL 100 mM CaCl_2_ was added for 30 mins on an ice-water bath to activate digestion. Next, digestion was quenched by adding 150 μL of 2X Low Salt STOP buffer (200 mM NaCl, 20 mM EDTA, 4 mM EGTA, 1% NP40, 0.2 mg/mL glycogen, and 0.05 ng/mL *S. cerevisiae* spike-in DNA) for 1 h at 4°C. The beads were then magnetized, and the supernatant was transferred to a fresh tube with 20 μL 5M NaCl and 1.5 μL RNase A (Invitrogen) for 20 min at 37°C. Then, 3 μL of 10% SDS and 2.5 μL of Proteinase K were added for 10 min at 70°C followed by phenol-chloroform-isoamyl alcohol purification, AMPure XP (Beckman Coulter) purification, and EtOH precipitation. Samples were resuspended in 50 μL 1XTE and stored at -20°C until library preparation. Libraries were prepared using an NEB Next Ultra II kit and paired end sequenced to a depth of ∼10,000,000 mapped reads/sample.

The integrity of sequencing data was verified by fastqc v0.11.9. Paired-end fastq files were aligned to the hg38 genome using bowtie2, PCR duplicates were filtered using Picard, and low-quality reads were removed with SAMtools as previously described (26–28). The reads were bioinformatically size selected for transcription factor size class (<120bp). deepTools was used to generate bigwigs and bigwig files were loaded onto the UCSC genome browser for visualization.

### FBXO24^-/-^ mice

FBXO24^-/-^ mice were generated at the Mouse Transgenic Core Facility at the University of Pittsburgh using CRISPR/Cas9 technology as described before (29). In brief, 50 ng/ul guide RNAs (5′ AATCTCGATAGCAGGGTATC and 3′ GGGATGGCGAATGAAATGCT) and 100 ng/uL CAS9 mRNA were injected into C57BL/6J fertilized embryos. Embryos were implanted into surrogate C57BL6/J mice, and a male offspring with 581 bp deletion in exon 5 of the FBXO24 gene (chromosome 5 coordinates 137622103-137621522) was selected. FBXO24^-/-^ mice were without obvious phenotypic abnormalities and were backcrossed seven times into the C57BL/6J strain.

### Mass Spectroscopy

BEAS-2B cells were seeded into twenty 10 cm dishes. After 24 h, FoxP1-V5 and HA-Ub vectors were transfected (0.5 ug/mL). 48 h after transfection, cells were treated with 25 μM MG132 for 1 hour to block the proteasome, then treated with 4% CSE for 2 h to enhance ubiquitination of FoxP1 protein, and protein lysate collected in RIPA buffer supplemented with protease-phosphatase inhibitors (1:100) and ubiquitin aldehyde to inhibit deubiquitinases (1:100). 8000 μg of the collected protein was immunoprecipitated with 5 μg anti-V5 antibody, and the immunoprecipitated proteins were separated by electrophoresis using a 10% SDS gel. After electrophoresis, the gel was stained with Coomassie blue and imaged.

Gel bands containing proteins of interest were excised, and proteins were digested with trypsin as previously described (18). Briefly, gel bands were diced into small pieces (<1 mm^3^) and washed with 50% acetonitrile/ 25 mM ammonium bicarbonate until no more visible stain was present. The gel pieces were then dehydrated with 100% acetonitrile (ACN), reduced with 10 mM dithiothreitol (DTT) at 56°C for 1 h, followed by alkylation with 55 mM Iodoacetamide (IAA) at room temperature for 45 min in the dark. Excess DTT and IAA were removed by washing the gel pieces with 25 mM ammonium bicarbonate and then twice with 100% ACN. A solution containing 20 ng/µL sequencing grade modified trypsin and 25 mM ammonium bicarbonate was added to cover the gel pieces, and digestion was carried out overnight at 37°C. The resultant tryptic peptides were extracted from the gel with 70% ACN/5% formic acid (FA), vacuum dried, and reconstituted in 18 µL 0.1% FA for nLC-MS/MS analysis.

Tryptic peptides were analyzed by nanoflow liquid-chromatography tandem mass spectrometry (nLC-MS/MS) using a NanoAcquity UPLC (Waters’ Corporation, Milford, MA) interfaced to a Velos Pro linear ion trap mass spectrometer (Thermo Fisher Scientific, Waltham, MA). For each analysis, a 1 µL volume of protein digest was injected onto a C18 column (PicoChip™column packed with Reprosil C18 3μm 120Å chromatography media in a 10.5cm long, 75μm ID column with a 15μm tip, New Objective, Inc., Woburn, MA) and then eluted off to the mass spectrometer using a 37-minute linear gradient of 3-35% ACN/0.1% FA at a flow rate of 300 nL/min. The Velos Pro was operated in positive ionization mode with a spray voltage of 1.95 kV and capillary temperature of 275°C. Acquisition consisted of cycles of one full-scan MS1 (AGC of 3×104, 75 ms maximum ion accumulation time, and m/z range of 375-1800) followed by eight MS/MS spectra recorded sequentially for the most abundant ions in the ion trap (minimum signal required 1000 counts, 1×104 AGC target, 100 ms maximum injection time, isolation width 2 m/z, normalized collision energy 35, and activation time 10 ms). Dynamic exclusion (30 s) was enabled to minimize redundant selection of peptides previously selected for MS/MS.

Collected MS/MS spectra were searched using the MASCOT search engine v2.4.0 (Matrix Science Ltd., London, England) (19) against a SwissProt Human protein sequence database using the following modifications: static modification of cysteine carbamidomethylation (+57), variable modifications of methionine oxidation (+16), protein N-terminal acetylation (+42), and di-glycine remnant on lysine (+114). The mass tolerance was set to 1.4 Da for the precursor ions and 0.8 Da for the fragment ions. Peptide identifications were filtered using the PeptideProphetTM and ProteinProphet® algorithms with a protein and peptide threshold cutoff of 1% false discovery rate (FDR), and a minimum of two peptides per protein implemented in ScaffoldTM v5.0.1 (Proteome software, Portland, OR).

### Cigarette smoke extract (CSE)

CSE solutions were prepared using a modification of the method of Blue and Janoff (30). 40 mL of cigarette smoke from a 1R6F research cigarette (University of Kentucky, Kentucky, USA) was drawn into a 50 mL syringe containing 10 mL DMEM and mixed with the medium by vigorous shaking. The process was repeated till the entire cigarette was consumed. The resulting solution was sterile filtered (0.22 μm) and designated as 100% CSE solution. The solution was prepared fresh before each experiment and used within 15 minutes.

### Cigarette smoke exposure of mice

Mice were exposed to cigarette smoke in a whole body exposure chamber (4 cigarettes/day, 5 days/week; Kentucky research cigarette 1R6F) or to room air for two weeks as described previously (31).

### Reagents

Details about the antibodies and primers used can be found in the online supplement. Nonsense (scramble/negative control) dsiRNA (dsiScramble, Cat # 51-01-19-09), human dsi FBXO24 (hs.Ri.FBXO24.13.3) and mouse dsi FBXO24 (mm.Ri.FBXO24.13.1), human dsi FoxP1 (hs.Ri.FOXP1.13.1) and human dsi CAND1 (hs.Ri.CAND1.13.1) were obtained from Integrated DNA Technologies (IDT). MG132 was obtained from Cell Signaling (#2194), Ubiquitin aldehyde from Enzo (#BMC-UW8450-0050), protein A/G agarose from Santa Cruz Biotechnology (#sc 2003), and Halt protease phosphatase inhibitor cocktail from Invitrogen (#1861279).

### Statistical Analysis

Descriptive statistics are reported as mean ± standard deviation (SD). Non-parametric statistical tests were performed for all analyses unless stated otherwise using STATA v13 (StataCorp, College Station, TX) and GraphPad Prism v7.0 (GraphPad Software, La Jolla, CA).

## RESULTS

### Cullin-associated and NEDD8-dissociated protein 1 (CAND1) binds to FoxP1 protein and regulates its cellular levels

We performed nLC-MS/MS to identify binding partners for ubiquitinated FoxP1 protein. In addition to FoxP1, 194 proteins were repeatedly identified in four independent nLC-MS/MS experiments (Supplemental Table 1). Protein-protein interaction analysis using Molecular Complex Detection (MCODE) algorithm (32, 33) identified expected protein networks for a transcription factor such as proteins involved in protein DNA complex assembly, DNA repair, and Condensin 1 complex (**Fig 1A**). Similarly, pathway analysis of identified proteins suggested that nucleic acid and nucleotide binding were among the top enriched pathways (**Fig 1B**).

**Figure 1:**
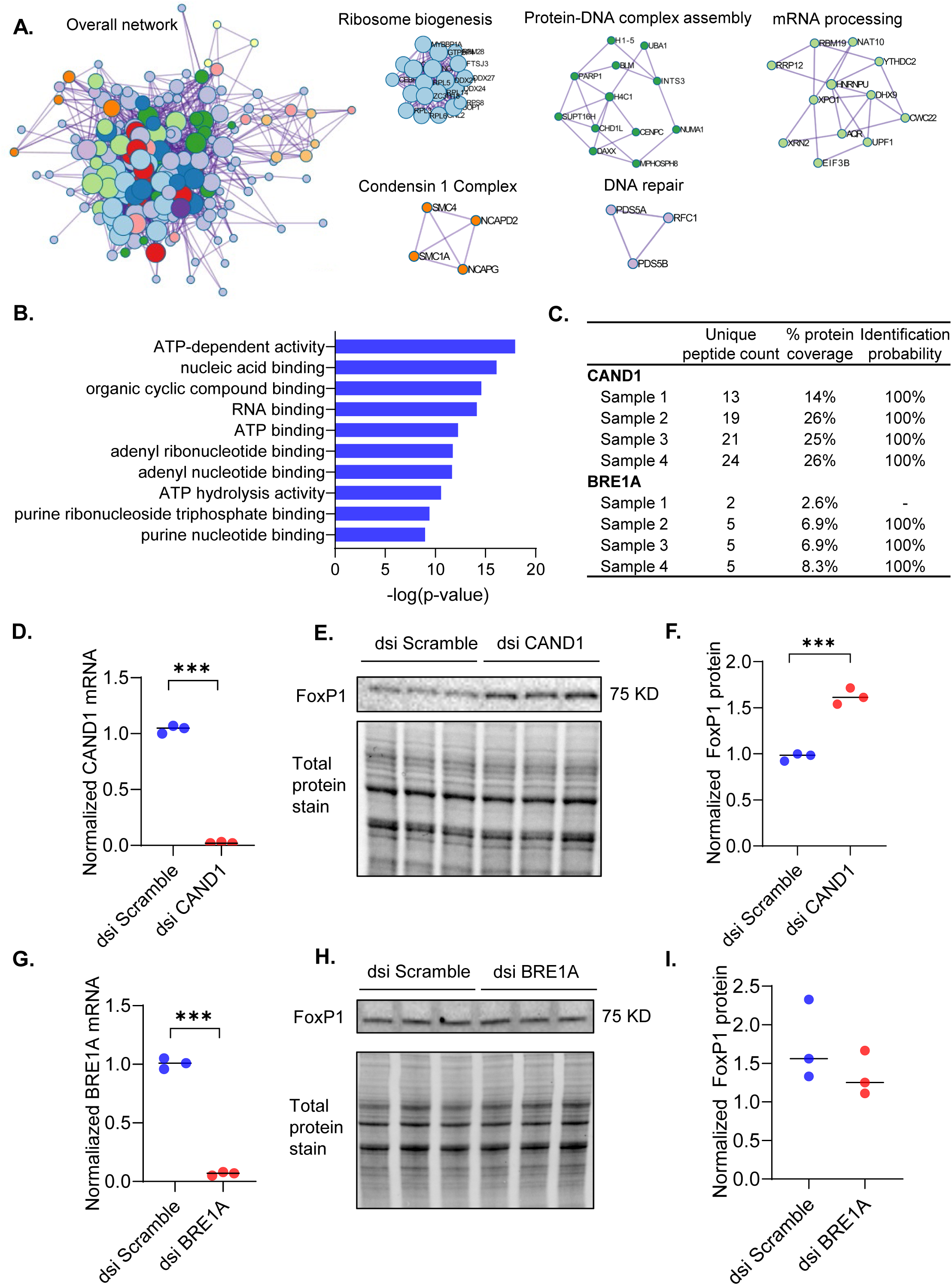
Cullin-associated NEDD8-dissociated protein 1 (CAND1) interacts with FoxP1 protein and regulates FoxP1 protein levels. FoxP1-V5 and HA-Ub were overexpressed in BEAS-2B cells. 48 h later, cells were incubated with 25 uM MG132 for 1 h to block the proteasome, followed by 4% CSE for 2 h to enhance FoxP1 protein ubiquitination. Cell lysate was then immunoprecipitated with anti-V5 antibody and eluate separated by gel electrophoresis. The gel was stained with Coomassie blue, and visible bands were excised and processed for mass spectroscopy by nLC-MS/MS. Protein-protein interaction analysis using Molecular Complex Detection (MCODE) algorithm **(A)** and pathway analysis **(B)** of 194 proteins consistently identified in 4 experiments. Peptide counts, protein coverage, and identification probability for CAND1 and BRE1A in the 4 experiments **(C)**. Knockdown of CAND1 followed by qPCR for CAND1 mRNA **(D)** and immunoblot for FoxP1 protein **(E-F)**. Knockdown of BRE1A followed by qPCR for BRE1A mRNA **(G)** and immunoblot for FoxP1 protein **(H-I)**. *** indicates *p*<0.001; *p*- values are from non-parametric tests from at least 3 independent experiments.

Three proteins involved in ubiquitination were identified. These included UBA1 (Ubiquitin Like Modifier Activating Enzyme 1), which is a ubiquitin E1 enzyme that regulates the stability of DNA repair proteins in the nucleus (34, 35). Also, Cullin-associated NEDD8-dissociated protein 1 (CAND1, **Fig 1C**) and the ubiquitin E3 ligase BRE1A were detected in all pull-downs. Spectral counts for CAND1 were higher than those for BRE1A (**Fig 1C**). While CAND1 is not a ubiquitin E3 ligase, it regulates the assembly and disassembly of Cullin-RING E3 ubiquitin ligase complexes (CRLs)(36, 37). BRE1A, also known as RNF20 (Ring Finger Protein 20), works in conjunction with another RING-type E3 ligase, BRE1B, to monoubiquitinate histone H2B at lysine 120 to regulate DNA transcription (38, 39). However, BRE1A is not known to ubiquitinate FoxP1 protein.

To determine if BRE1A or CAND1 can regulate FoxP1 protein levels, we assessed the impact of depleting BRE1A or CAND1 mRNA by RNAi on FoxP1 protein levels in lung epithelial cells. While the knockdown of CAND1 robustly increased FoxP1 protein levels (**Fig 1D-F**), knockdown of BRE1A had no effect (**Fig 1G-I**).

These data suggest that CAND1 binds to and regulates FoxP1 protein levels and, therefore, a Cullin-RING ubiquitin E3 ligase may regulate FoxP1 protein stability.

### The Skp-Cullin-F-box (SCF) family E3 ligase FBXO24 regulates cellular FoxP1 protein levels at the post-translational level

Among the Cullin-RING E3 ligases, the Skp-Cullin-F-box (SCF) E3 ligases form the largest subfamily (40) and are the most extensively studied in terms of their interactions with CAND1 (36, 37). Accordingly, we overexpressed a library of SCF E3 ligases in lung epithelial cells to identify the E3 ligase that may regulate FoxP1 protein levels. The only SCF family E3 ligase that reduced FoxP1 protein levels in a dose-dependent manner upon overexpression was FBXO24 (**Fig 2A-B**), while other E3 ligases, such as FBXO23, did not (**Fig 2C**).To confirm, we overexpressed a murine FBXO24 construct in mouse lung epithelial cells and again noted a reduction in FoxP1 protein levels (**Fig 2D**). For further confirmation, we knocked down FBXO24 by RNAi in BEAS-2B cells and MLE-12 cells and noted that FoxP1 protein levels significantly increased (**Fig 2E-G**). Knockdown of FBXO24 increased FoxP1 protein levels without altering FoxP1 mRNA levels, suggesting that FBXO24 regulates FoxP1 post-translationally, as would be expected for an E3 ligase (**Fig 2H**). These data suggested that the Skp-Cullin-F-box (SCF) family E3 ligase FBXO24 robustly regulates cellular FoxP1 protein levels at the post-translational level.

**Figure 2:**
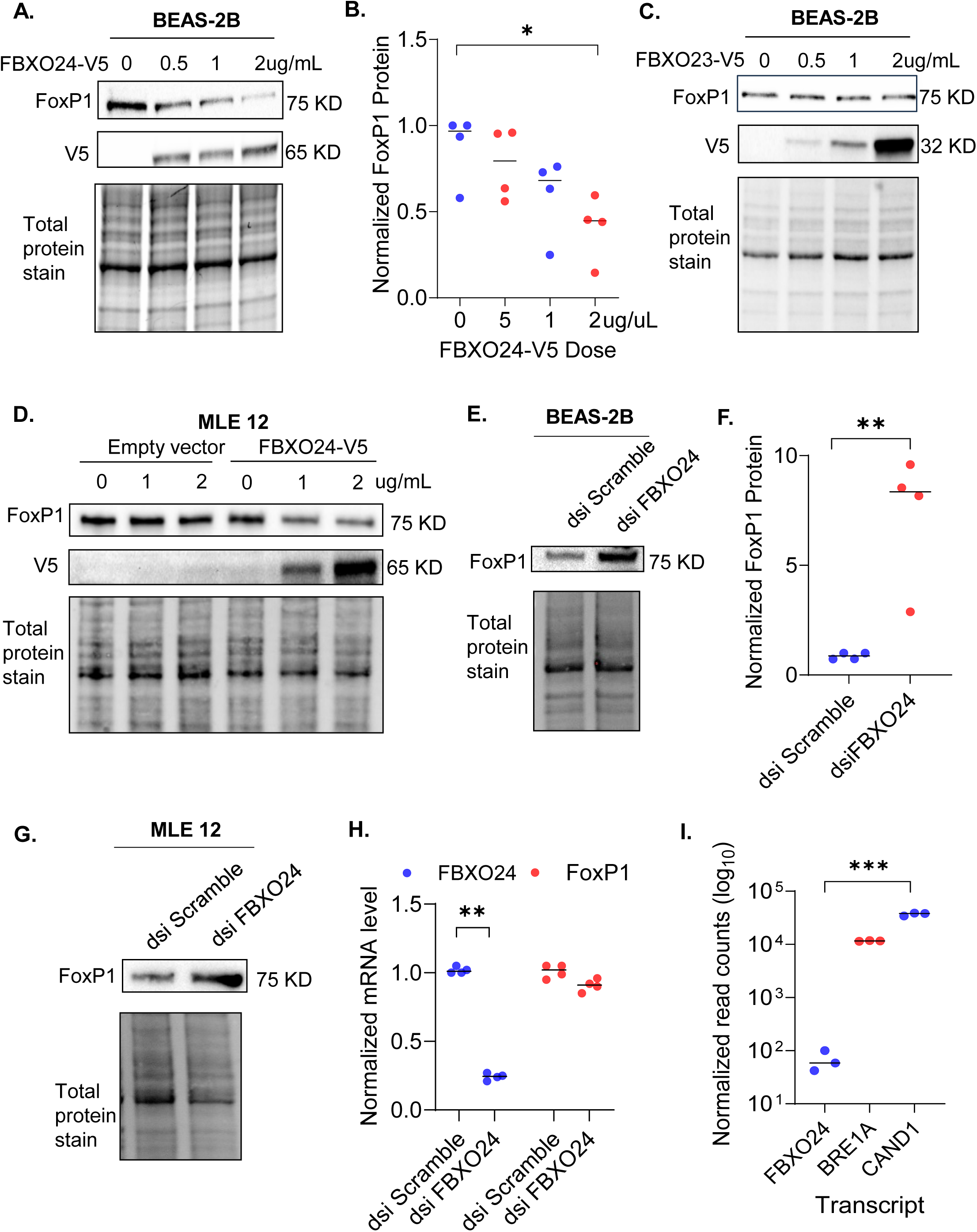
The E3 ligase FBXO24 regulates FoxP1 protein levels. Increasing concentrations of FBXO24-V5 plasmid were transfected into BEAS-2B cells, followed by immunoblotting for FoxP1 and V5 24 h later **(A-B)**. Increasing concentrations of FBXO23-V5 plasmid were transfected into BEAS-2B cells, followed by immunoblotting for FoxP1 and V5 24 h later **(C)**. Increasing concentrations of FBXO24-V5 plasmid or vector backbone were transfected into MLE 12 cells, followed by immunoblotting for FoxP1 and V5 24 h later **(D)**. FBXO24 was depleted in BEAS-2B cells by transfecting 30 nM dicer-substrate short interfering RNAs targeting FBXO24 (dsi FBXO24) vs. dsi Scramble (control) for 48 h, followed by immunoblotting for FoxP1 **(E-F)**. FBXO24 was depleted in MLE-12 cells by transfecting 30 nM dsi FBXO24 vs. dsi Scramble (control) for 48 h, followed by immunoblotting for FoxP1 and qPCR for FBXO24 and FoxP1 mRNA **(G-H)**. Normalized read counts for FBXO24, CAND1, and BRE1A mRNA in RNA-seq data from BEAS-2B cells **(I)**. * indicates *p*<0.05, ** indicates *p*<0.01; *** indicates *p*<0.001. *p*-values are from nonparametric tests from 4 independent experiments.

Next, we examined bulk RNA sequencing data from BEAS-2B cells (11) to determine why FBXO24 was not identified in our mass spectroscopy experiments. We found that FBXO24 transcript levels (<100 reads/cell) were more than 100-fold less than CAND1 and BRE1A transcript levels (**Fig 2I**). The low transcript numbers for FBXO24 suggested that FBXO24 protein was present in low levels in the cell, making it difficult to identify by mass spectroscopy, unlike CAND1 and BRE1A protein, which were likely much more abundant.

### FBXO24 protein complexes with and colocalizes to the nucleus with FoxP1 protein and regulates K48 and K63 ubiquitination of FoxP1 protein

To determine if FBXO24 protein complexes with FoxP1 protein, we performed co-immunoprecipitation and immunostained lung epithelial cells for both proteins. Results suggested that FBXO24 co-immunoprecipitates with FoxP1 protein (**Fig 3A**). Also, FBXO24 protein was found to colocalize to the nucleus with FoxP1 protein (**Fig 3B**).

**Figure 3:**
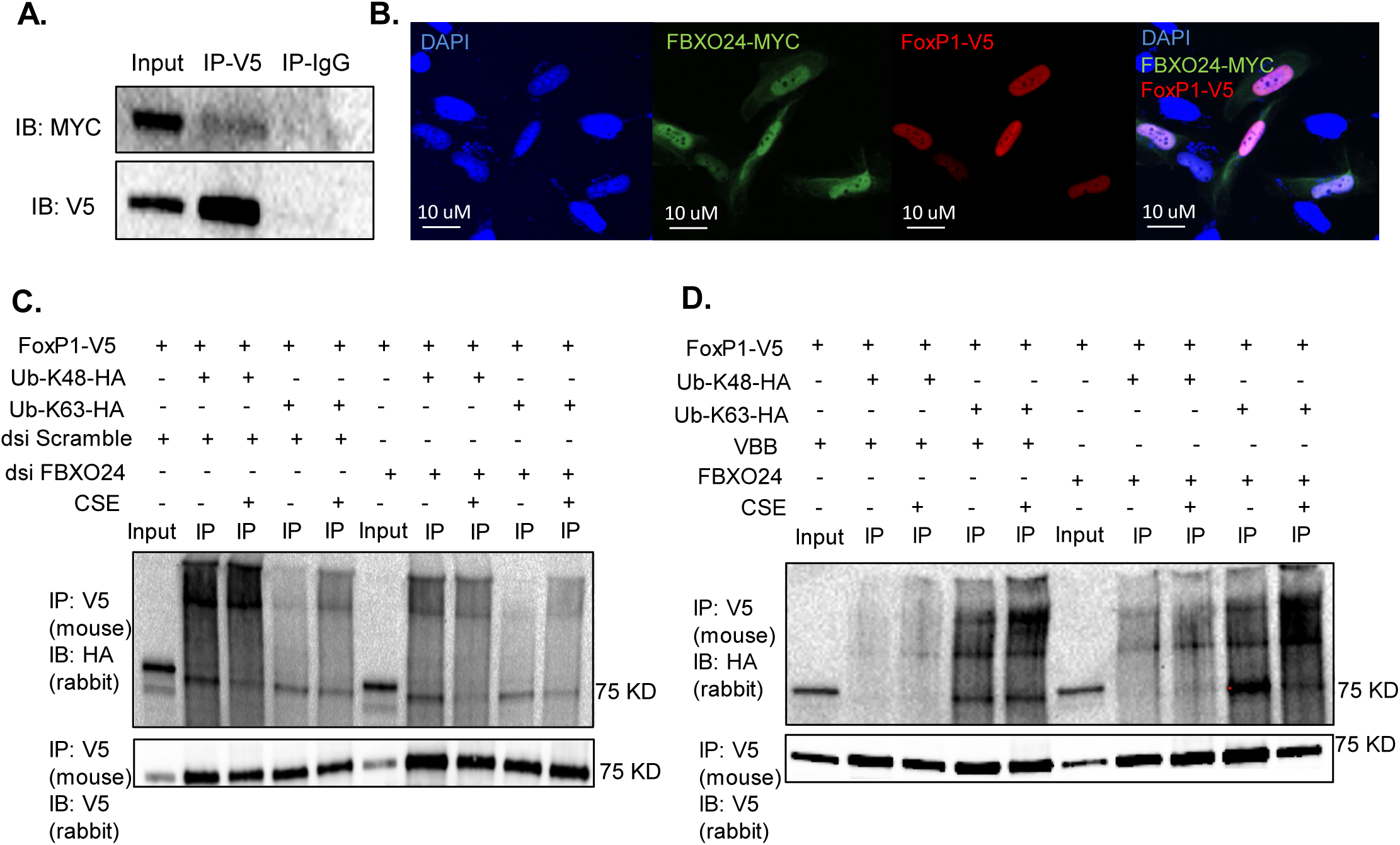
FBXO24 protein complexes with and colocalizes to the nucleus with FoxP1 protein and regulates K48 and K63 ubiquitination of FoxP1 protein. FoxP1-V5 and FBXO24-MYC plasmids were transfected into BEAS-2B cells for 24 h. Cell lysate was then immunoprecipitated with mouse anti-V5 antibody or normal mouse IgG and immunoblotted with rabbit anti-MYC and rabbit anti-V5 antibodies **(A)**. FoxP1-V5 and FBXO24-MYC plasmids were transfected into BEAS-2B cells for 24 h. Cells were then immunostained with anti-MYC antibody (green) and anti-V5 antibody (red) along with DAPI (blue) **(B)**. 1 ug/mL of FoxP1-V5 plasmid was transfected into BEAS-2B cells, along with either HA-tagged mutated ubiquitin that can only form K48 chains (Ub-K48-HA) or K63 chains (Ub-K63-HA) with or without knockdown of FBXO24 by RNAi. Cells were incubated with 25 uM MG132 for 1 h and then exposed to 2% CSE for 2 h. Cell lysate was immunoprecipitated with mouse anti-V5 antibody and immunoblotted with rabbit anti-HA **(C, top)** and rabbit anti-V5 antibodies **(C, bottom)**. FoxP1-V5 plasmid was transfected into BEAS-2B cells, along with either HA-tagged mutated ubiquitin that can only form K48 chains (Ub-K48-HA) or K63 chains (Ub-K63-HA) with or without overexpression of FBXO24. Cells were incubated with 25 uM MG132 for 1 h and then exposed to 2% CSE for 2 h. Cell lysate was immunoprecipitated with mouse anti-V5 antibody and immunoblotted with rabbit anti-HA **(D, top)** and rabbit anti-V5 antibodies **(D, bottom)**.

Our prior work demonstrates that FoxP1 protein undergoes K48 and K63 ubiquitination, and exposure of lung epithelial cells to cigarette smoke extract increases both kinds of ubiquitination (11). To confirm that FBXO24 regulates FoxP1 protein ubiquitination, we examined K48 and K63 ubiquitination of FoxP1 protein with overexpression and knockdown of FBXO24 in lung epithelial cells. Results suggested that the knockdown of FBXO24 reduced K48 and K63 ubiquitination of FoxP1 protein with and without exposure to cigarette smoke extract (**Fig 3C**). In agreement, overexpression of FBXO24 increased K48 and K63 ubiquitination of FoxP1 protein with and without exposure to cigarette smoke extract (**Fig 3D**).

These studies suggest that FBXO24 binds to FoxP1 protein in the nucleus and regulates its K48 and K63 ubiquitination.

### Knockdown of FBXO24 can reverse adverse changes in cellular function and viability resulting from loss of FoxP1 protein

The data above suggested that knocking down FBXO24 can increase FoxP1 protein levels; however, it remained unclear if the rescued FoxP1 protein was functional. Our previous research has shown that loss of FoxP1 protein leads to dysregulated overactivity of the UPR (11). Therefore, we investigated the UPR activity in lung epithelial cells after knocking down FBXO24 protein.

Results suggested that knockdown of FBXO24 reduced levels of various markers of UPR activity, such as transcript levels for the main protein folding chaperone in the endoplasmic reticulum, i.e., glucose-regulated protein 78 (GRP78, **Fig 4A**). In agreement, transcript levels for total and spliced (activated) XBP-1, PERK, and phosphorylation of EIF-2α (a marker of PERK activation, **Fig 4B-E**) were reduced in the control condition and in the presence of cigarette smoke extract (positive control). Finally, levels of the death mediator CHOP, that increase when the UPR is overactive, were also significantly reduced in the presence or absence of cigarette smoke extract (**Fig 4F**). In agreement, knockdown of FBXO24 reversed the loss of cell viability that occurred upon depletion of FoxP1 (**Fig 4G-I**).

**Figure 4:**
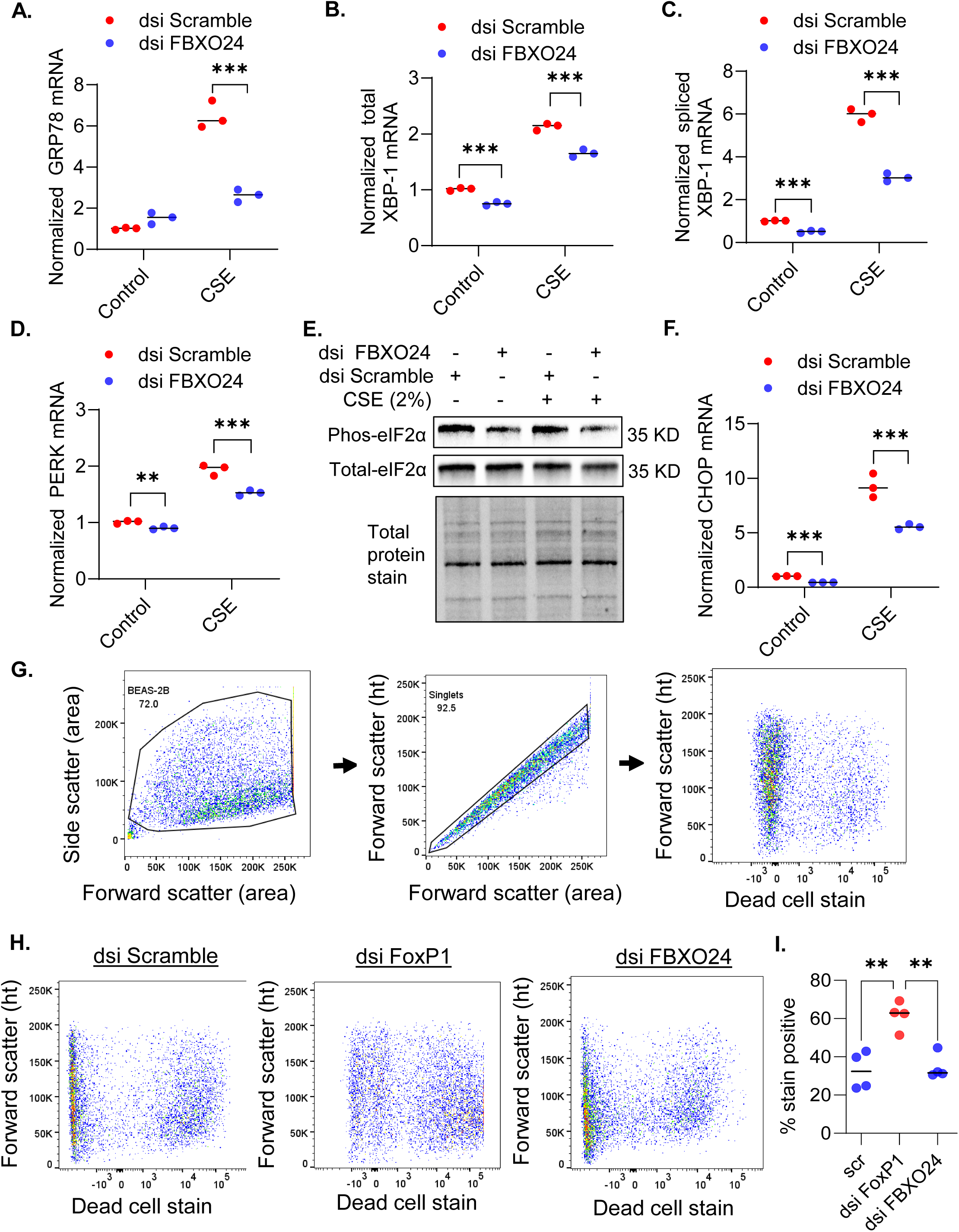
Knockdown of FBXO24 reduced activity of the unfolded protein response. FBXO24 was depleted in BEAS-2B cells by transfecting 30 nM dicer-substrate short interfering RNAs targeting FBXO24 (dsi FBXO24) vs. dsi Scramble (control) for 48 h with and without exposure to 2% cigarette smoke extract for 6 h to induce the UPR. Cell lysate was processed for qPCR for UPR mediators GRP78, total XBP-1, spliced XBP-1, PERK, and CHOP, and immunoblotting for total and phos-EIF2 alpha **(A-F)**. BEAS-2B cells were transfected with 30 nM dsi Scramble (control/scr), dsi FoxP1, or dsi FBXO24 for 48 h followed by flow cytometry (gating strategy per **G**) to determine cell viability using a vital dye **(H-I)**. *indicates *p*<0.05, **indicates *p*<0.01, & ***indicates *p*<0.001; *p* values are from nonparametric tests from 3 independent experiments.

These data suggest that suppressing FBXO24 can increase FoxP1 protein levels and reverse the harmful downstream consequences of reduced FoxP1 protein levels. Consequently, targeting FBXO24 may offer a promising therapeutic strategy for cardiovascular and pulmonary diseases characterized by reduced FoxP1 protein levels.

### Inhibiting FBXO24 increases FoxP1 protein levels and reduces cigarette smoke induced UPR activity in the lung *in vivo*

We sought to determine if inhibiting FBXO24 can reverse the adverse effects of reduced FoxP1 protein levels *in vivo*. To test our hypothesis, we induced the UPR in the lungs of FBXO24^-/-^ mice by exposing them to cigarette smoke for 2 weeks. As expected, FBXO24^-/-^ mice had increased FoxP1 protein levels in the lungs despite similar FoxP1 mRNA levels as wild-type littermates (**Fig 5A-E**). FBXO24^-/-^ mutants had less UPR activity in the lungs after exposure to cigarette smoke than control mice (**Fig 5F-H**). This finding may be significant because UPR activity in the lung contributes to common lung diseases such as COPD and pulmonary fibrosis (41, 42). Similarly, FBXO24^-/-^ mice had increased levels of FoxP1 protein in cardiac tissue despite similar FoxP1 mRNA levels as wild-type littermates (**Fig 5I-K**).

**Figure 5:**
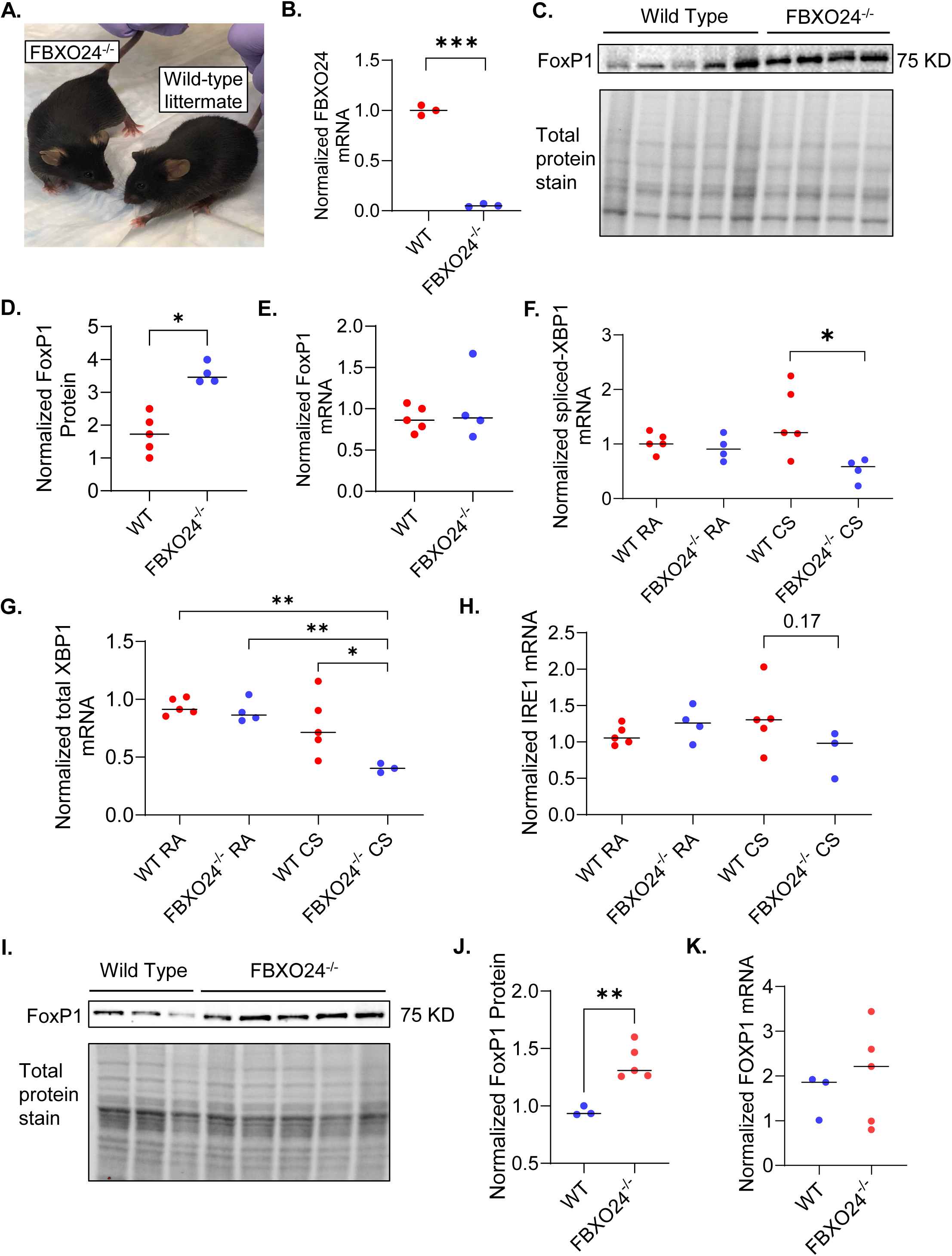
FBXO24^-/-^ mice have higher FoxP1 protein levels in the lungs and the heart and reduced UPR activity in the lungs after short-term exposure to cigarette smoke. FBXO24^-/-^ mice are viable without evident phenotypic abnormalities **(A)**. FBXO24 mRNA levels were assessed by qPCR in whole lung lysates from FBXO24^-/-^ mice and wild-type littermates **(B)**. FoxP1 protein levels by immunoblot **(C-D)** and FoxP1 mRNA levels by qPCR **(E)** in whole lung lysates from FBXO24^-/-^ mice and wild-type littermates. FBXO24^-/-^ mice and wild-type littermates were exposed to cigarette smoke for 2 weeks to induce the UPR or to room air (control) followed by qPCR to assess levels of spliced XBP-1 **(F)**, total XBP-1 **(G)**, and IRE1 mRNA **(H)** in whole lung lysates. FoxP1 protein levels by immunoblot **(I-J)** and FoxP1 mRNA levels by qPCR **(K)** in cardiac lysates from FBXO24^-/-^ mice vs. wild-type littermates. **\*** indicates *p*<0.05; **\*\*** indicates *p*<0.01.

### FoxP1 protein binds to and regulates the FBXO24 gene promoter

We found evidence of reciprocal regulation between FoxP1 and FBXO24. Specifically, knockdown of FoxP1 increased FBXO24 transcript levels (**Fig 6A**). Because FoxP1 is a transcriptional repressor, we computationally cross-referenced the DNA motif recognized by FoxP1 (**Fig 6B**) against the sequence of the FBXO24 gene promoter and found two high-probability binding sites within the promoter (**Fig 6C**). Also, we examined publicly available FoxP1 ChIP-seq data and found enrichment for FoxP1 over the FBXO24 promoter region in HepG2 cells (GEO: GSE105268) and embryonic stem cells (GEO: GSE30992, **Fig 6D**).

**Figure 6:**
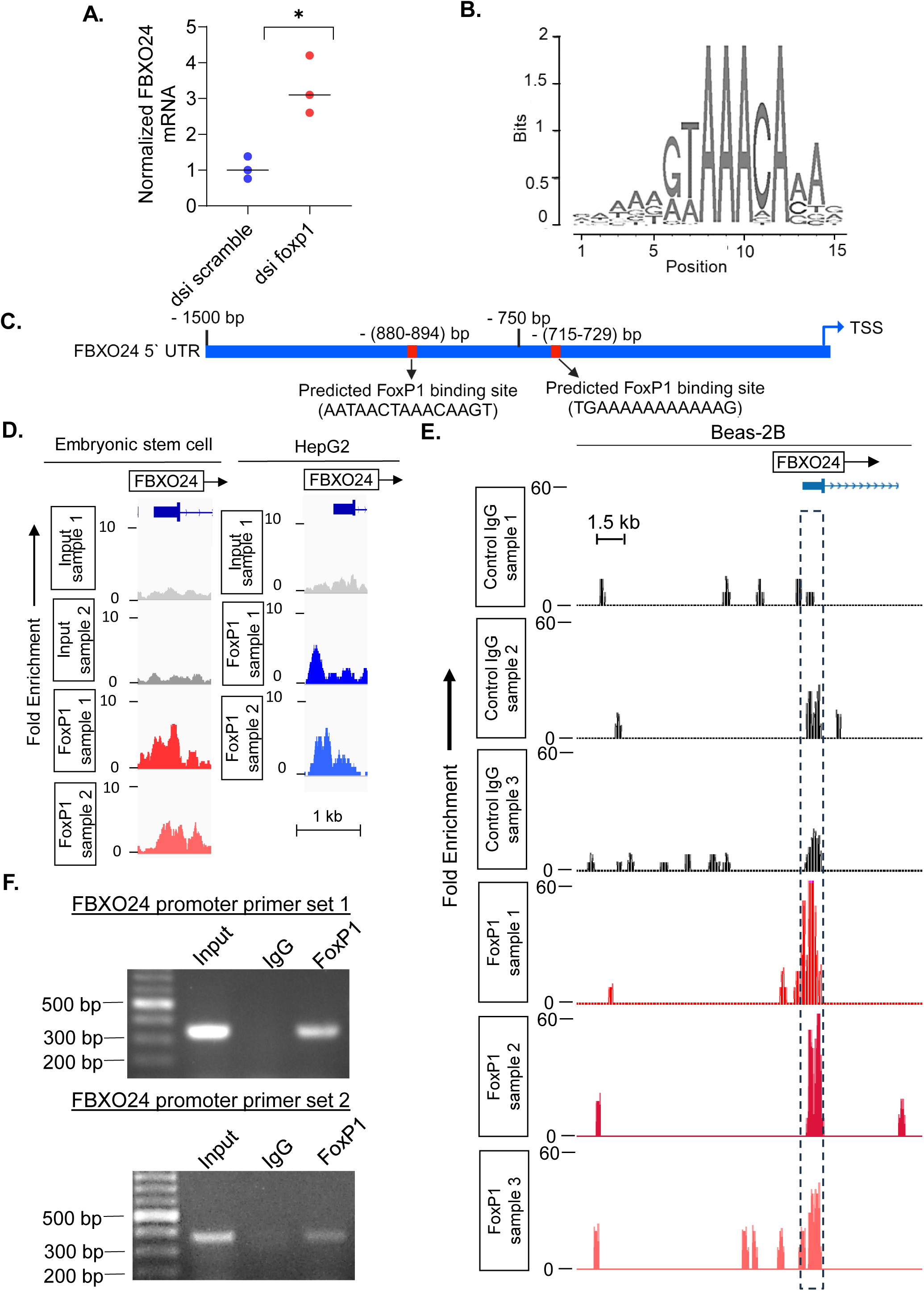
FoxP1 binds to and represses the FBXO24 promoter. Lung epithelial cells were transfected with 30 nM dsi RNA targeting nonsense mRNA (dsi Scramble) or FoxP1 mRNA. 48h later, cell lysate was subject to RT-PCR to assess FBXO24 mRNA levels **(A)**. The published probability matrix for the DNA motif recognized by FoxP1 protein (19) **(B)**. Computational analysis identified two high-probability binding sites in the 5′ UTR of the FBXO24 gene that matched the DNA motif recognized by FoxP1 protein (as depicted in panel B) **(C)**. Analysis of publicly available FoxP1 ChIP-Seq data from embryonic stem cells (GEO: GSE30992) and HepG2 cells (GEO: GSE105268) with browser tracks of the FBXO24 promoter region **(D)**. FoxP1-V5 was overexpressed in A549 cells. 48 h later, nuclei were isolated and subject to CUT&RUN using anti-V5 or control IgG. Browser tracks at the FBXO24 promoter region (enclosed in dotted box) were generated **(E)**. Endogenous FoxP1 ChIP in A549 cells was followed by PCR for the FBXO24 promoter region using two different sets of primers (primer sequence per online supplement). Amplicons were run on an agarose gel **(F)**. \**p*<0.05. Non-parametric tests were used (Mann-Whitney U).

To confirm these findings in lung epithelial cells, we performed CUT&RUN in A549 cells after overexpressing V5-tagged FoxP1 protein. Results suggested close to 60-fold enrichment for FoxP1 binding in the FBXO24 promoter (**Fig 6E**). Finally, to confirm that endogenous FoxP1 protein also binds to the FBXO24 promoter, we performed ChIP with a FoxP1 antibody in A549 cells followed by PCR for the FBXO24 promoter sequence. Results suggested that FoxP1 ChIP samples were enriched for the FBXO24 promoter sequence, unlike the IgG ChIP samples (control, **Fig 6F**).

These data suggest a positive feedback loop between FoxP1 and FBXO24, whereby loss of FoxP1 increases expression of FBXO24 that may further reduce FoxP1 protein levels. To our knowledge, this is the first example of a substrate regulating its E3 ligase and may have a number of implications.

## DISCUSSION

FoxP1 protein’s ability to mitigate a variety of prevalent cardiopulmonary disorders makes it a compelling target for further investigation. Specifically, recent studies report that reduced FoxP1 protein levels promote vascular inflammation in atherosclerosis (6), cardiac remodeling and fibrosis in heart failure (8, 9), and proteostatic stress due to tobacco consumption in COPD (10, 11). The present study identifies FBXO24 as the E3 ligase that controls FoxP1 protein levels. Specifically, we found that (1) CAND1, an adapter protein used by Cullin-RING E3 ubiquitin ligases, binds to and regulates FoxP1 protein levels using unbiased methods; (2) FBXO24 regulates cellular FoxP1 protein levels by modulating its K48 and K63 ubiquitination and complexing with FoxP1 protein in the nucleus; (3) inhibiting FBXO24 can rescue FoxP1 protein and reverse cellular dysfunction and reduced viability resulting from loss of FoxP1 protein *in vitro*; (4) inhibiting FBXO24 increases FoxP1 protein levels in the lung and heart and reduces cigarette smoke-induced proteostatic stress in the lung *in vivo*; and (5) FoxP1 binds to and represses the FBXO24 promoter to reduce FBXO24 transcript levels in the cell. While more proof-of-concept studies are needed, targeting FBXO24 might serve as a promising therapeutic strategy for common cardiopulmonary disorders. Small molecule FBXO24 inhibitors are already being developed (29).

FBXO24 is a relatively understudied E3 ligase, with only 10 publications mentioning it to date (29, 43–51). Three of these studies identified FBXO24 through high-throughput screens but did not investigate it further (46, 50, 51). Another two focused on how FBXO24 may regulate male fertility (48, 49). The remaining studies identified five FBXO24 substrates—NDKA (44), PRMT6 (43), LSD1 (44), FoxK2 (47), and DARS2 (29)—each linked to distinct biological processes.

Three of the five substrates ubiquitinated by FBXO24 regulate carcinogenesis. First, Nucleoside Diphosphate Kinase A (NDKA), responsible for synthesizing nucleoside triphosphates other than ATP (52), has reduced expression levels in metastatic cancer cells (53, 54). Also, mutations in NME1, the gene encoding NDKA, have been identified in patients with aggressive neuroblastomas, highlighting NDKA’s tumor-suppressive properties (55). Second, Protein Arginine Methyltransferase 6 (PRMT6), which functions as a coregulator of gene expression through methylation of histone residues H3R2, H4R3, and H2AR3, is aberrantly expressed in various cancers (56, 57). PRMT6 is an oncogenic modifier that can both repress or promote the progression of cancer but does not initiate cancer in healthy tissues (58, 59). Finally, Lysine-specific demethylase 1 (LSD1), a crucial chromatin modulator responsible for demethylating H3K4me1/2, is involved in tumorigenesis. FBXO24 negatively regulates LSD1, thereby suppressing tumorigenesis in breast cancer cells (45).

Further, FBXO24 has been reported to ubiquitinate two important proteins during bacterial infection, Forkhead box protein K2 (FoxK2) and Aspartyl-tRNA synthetase (DARS2)(29, 47). Specifically, FBXO24 ubiquitinates and degrades FoxK2 in lung epithelial cells infected with *Pseudomonas aeruginosa* or *Klebsiella pneumoniae* (47). The resulting loss of FoxK2 impairs mitochondrial function in these cells. In agreement, FBXO24^-/+^ mice exhibit preserved mitochondrial function and higher FoxK2 protein levels in their lungs compared to wild-type littermates during bacterial pneumonia (47). Also, FBXO24 ubiquitinates and degrades DARS2 in lung epithelial cells during bacterial pneumonia (29). DARS2 is a mitochondrially derived enzyme released into the circulation during these infections that possesses intrinsic innate immune and cellular repair properties (29). Consistent with these findings, FBXO24^-/-^ mice displayed increased pulmonary levels of DARS2 and enhanced cellular and cytokine responses during experimental pneumonia (29). Therefore, inhibiting FBXO24 may promote anti-pathogen immune responses and preserve mitochondrial function in lung epithelia, promoting the host’s ability to overcome bacterial infections (60).

These studies suggest that inhibiting FBXO24 will have diverse and context-dependent biological consequences. Specifically, inhibition of FBXO24 would concurrently elevate levels of a tumor suppressor (44), an oncogenic modifier (58), and a cancer-promoting protein (58), necessitating careful evaluation of the overall impact. Our (preliminary) observations in FBXO24^-/-^ mice revealed no increase in the incidence of spontaneous tumors. Also, inhibiting FBXO24 should increase host defense against bacterial lung infections (29, 47). However, it may also impair male fertility (48, 49), as evidenced by infertility in FBXO24^-/-^ male mice observed in our colony, while FBXO24^-/+^ male and FBXO24^-/-^ female mice bred normally. These observations are consistent with prior reports that FBXO24 is required for normal spermatogenesis (48, 49). Finally, inhibiting FBXO24 should reduce cigarette smoke-induced UPR activity in COPD, pathological cardiac remodeling in heart failure, and vascular inflammation in atherosclerosis. Nonetheless, successful clinical translation will require careful consideration of context-specific outcomes of stabilizing various substrate proteins.

We are the first to report that a substrate of an E3 ligase can reciprocally regulate its E3 ligase, a finding with several significant implications. First, it indicates that one substrate of an E3 ligase can indirectly control the degradation of other substrates through modulation of the ligase, revealing previously unrecognized regulatory networks. Specifically, in the context of FoxP1, a reduction in FoxP1 protein levels may enhance FBXO24 activity, subsequently lowering the levels of other FBXO24 substrates such as FoxK2, DARS2, and NDKA. The loss of these other substrates may contribute to the increased susceptibility to lung infections and cancer observed in COPD patients, who characteristically exhibit reduced FoxP1 protein levels in the lungs. Second, therapeutic inhibition of FBXO24 would increase FoxP1 protein levels, thereby further decreasing FBXO24 transcript levels. Thus, even moderate inhibition of FBXO24 could result in substantial therapeutic benefits due to the amplification effect from increased FoxP1 protein levels. Lastly, this positive feedback loop, where increased FoxP1 protein levels suppress FBXO24 mRNA levels, provides a plausible explanation for why FBXO24 transcript levels are exceptionally low under steady-state conditions (Fig. 2I). Further elucidation of these regulatory mechanisms will be essential for developing targeted and effective therapeutic strategies involving FBXO24.

Our study has limitations. First, we had trouble detecting endogenous FBXO24 protein by mass spectrometry or immunoblot in lung epithelial cells, likely due to the very low level of expression of FBXO24 (Fig 2I). Also, further work is necessary to fully elucidate the structural basis of the interaction between FBXO24 and FoxP1. Our preliminary findings indicate that FoxP1 protein undergoes multisite ubiquitination with different types of ubiquitin linkages, suggesting complex interaction with its E3 ligase. Finally, we demonstrated that FBXO24^-/-^ mice exhibit elevated FoxP1 protein levels in the lungs and the heart, along with reduced UPR activity in the lungs following 2 week cigarette smoke exposure. However, long-term studies with tissue-specific FBXO24 knockout models will be needed to demonstrate that these observations translate into protection against cardiopulmonary disease.

This study identifies FBXO24 as the ubiquitin E3 ligase that controls FoxP1 protein stability by regulating its K48 and K63 ubiquitination. FoxP1 protein levels are reduced in prevalent cardiopulmonary diseases such as COPD, atherosclerosis, and heart failure, where they causally contribute to disease pathophysiology. Our findings suggest that inhibiting FBXO24 can increase FoxP1 protein levels in the lungs and the heart, highlighting the therapeutic potential of FBXO24 inhibition. Notably, we uncover reciprocal regulation where FoxP1 protein modulates FBXO24 expression, revealing new regulatory networks in ubiquitin-proteasome biology.

### SOURCES OF FUNDING

This project was supported by R01HL153400 to DC. This project was supported by UPMC Hillman Cancer Center Proteomics Facility supported in part by award P30CA047904. This research was supported in part by the University of Pittsburgh Center for Research Computing, RRID:SCR_022735, through the resources provided. Specifically, this work used the HTC cluster, which is supported by NIH award number S10OD028483. This project used the University of Pittsburgh HSCRF Genomics Research Core, RRID: SCR_018301 NGS sequencing services, with special thanks to the Assistant Director, Will MacDonald. This work was also supported by National Institutes of Health R01HL153400, K23HL126912 and R35 GM133732 to S.J.H and by P01HL114453, R01HL097376, R01HL081784, and R01HL096376 awarded to RKM.

## Abbreviations

Fbx: F-box
SCF: Skp-Cullin-F box
UPS: ubiquitin-proteasome system
FoxP1: Forkhead Box Protein P1
UPR: unfolded protein response

