## Supplemental Materials for "Reciprocal Regulation Between the SCF^FBXO24^ Ubiquitin E3 Ligase and FoxP1 Protein"

**Human Primer sequences for RT-PCR:**

|  | Species | FWD | REV |
| --- | --- | --- | --- |
| GAPDH | Human | TCATTTCCTGGTATGACAACGA | GTCTTACTCCTTGGAGGCC |
| FoxP1 | Human | TGCTCAAGGCATGATTCCA | CCTGTGGTTTCTTCTGCAG |
| FBXO24 | Human | CCTTGGCCTGGTGGATGAAT | TCGGTCCATTTTGTCCCCTG |
| GRP78 | Human | GGATCATCAACGAGCCTACG | CACCCAGGTCAAACACCAG |
| CHOP | Human | AAGGCACTGAGCGTATCATGT | TGAAGATACACTTCCTTCTTGAACA |
| Total XBP-1 | Human | TTACGAGAGAAAACTCATGGCC | GGGTCCAAGTTGTCCAGAATGC |
| Spliced XBP-1 | Human | CTGAGTCCGAATCAGGTGCAG | ATCCATGGGGAGATGTTCTGG |
| PERK | Human | GTCCGGAACCAGACGATGAG | GGCTGGATGACACCAAGGAA |

**Murine Primer sequences for RT-PCR:**

|  | Species | FWD | REV |
| --- | --- | --- | --- |
| GAPDH | Mouse | AACTTTGGCATTGTGGAAGG | ACACATTGGGGGTAGGAACA |
| FoxP1 | Mouse | GTTAGAGCTACAGCTTGCA | TTGGGTTCTGTAGACTTCAC |
| FBXO24 | Mouse | TACGTGGTGTTGTGTCGAGG | GACCTCCACACAGTCACAGG |
| CHOP | Mouse | AACAGAGGTCACACGCACAT | ACTTTCCGCTCGTTCTCCTG |
| Spliced XBP-1 | Mouse | CTGAGTCCGAATCAGGTGCAG | GTCCATGGGAAGATGTTCTGG |
| PERK | Mouse | TCTTGGTTGGGTCTGATGAAT | GATGTTCTTGCTGTAGTGGGGG |
| IL-6 | Mouse | TACCACTTCACAAGTCGGAGGC | CTGCAAGTGCATCATCGTTGTTC |

**Human Primer sequences for ChIP-PCR:**

|  | FWD | REV |
| --- | --- | --- |
| FBXO24 set 1 | GTCCCCCAAAGACCAATCGT | CAGAGGCTTCACAGTCGGAG |
| FBXO24 set 2 | AGGCTGCAGTCTCCTTTTGG | AGCTCCTGGGTTCTAGCTCA |

**Antibodies**

|  | Host species | Application | Cat No |
| --- | --- | --- | --- |
| FoxP1 | Mouse | WB | sc-398811, Santa Cruz |
| FoxP1 | Rabbit | WB | ab16645, Abcam |
| V5 | Mouse | WB, IF, IP | SV5-Pk1, Bio-Rad |
| V5 | Rabbit | WB, IF, IP | # 13202, Cell signaling |
| MYC | Rabbit | WB, IF | # 05724, Millipore |
| HA | Rabbit | WB | # 3724, Cell signaling |
| Total-eIF2α | Rabbit | WB | # 5324, Cell signaling |
| Phos-eIF2α | Rabbit | WB | # 3398, Cell signaling |
| Anti-mouse secondary | Goat | WB, IF | 1706516, Bio Rad |
| Anti-rabbit secondary | Goat | WB, IF | 111-035003, Jackson |
